# Conspecific attraction in songbirds during spring migration

**DOI:** 10.1101/2025.10.20.683507

**Authors:** André Desrochers, Laetitia Desbordes

## Abstract

Conspecific attraction has often been hypothesized as a mechanism that could facilitate migration and territory acquisition. We studied conspecific attraction by songbirds during spring migration between 2019 and 2025 at the Dunes of Tadoussac, Quebec, Canada. This location is known to produce spectacular migration events with thousands of passerine birds flying at close range from the observers. We recorded sequences of individual birds moving just over the ground in a ∼20 m wide corridor along the edge of the St. Lawrence estuary. We also conducted hourly counts to detect conspecific clusters at a coarser temporal scale. Birds were much more likely to be following conspecifics at close range than other species, as evidenced by randomization tests, after accounting for species composition and abundance during each specific migration event. Conspecific sequences were mostly of two individuals, but much larger sequences occurred, up to 103 consecutive individuals of the same species. We found no evidence of leading species, i.e. species that were more likely to lead, rather than follow, another species. Hourly species counts were often much more variable than expected from a random (Poisson) process, at least in the case of Tennessee and Magnolia Warblers, providing evidence for conspecific attraction at a coarser temporal scale. This study provides the first detailed evidence of conspecific attraction with passerine birds during migration.

## 1 Introduction

Bird migrations often assemble birds of different species in stopover sites and along migration corridors (Gauthreaux 1972; Cohen and Satterfield 2020). This phenomenon, termed *co-migration*, fosters interspecific interactions, leading to nonrandom species assemblages (Van Doren et al. 2025; DeSimone et al. 2024). Those interspecific interactions may be agonistic (Recher and Recher 1969; Novcic 2018), occasionally lethal (Ydenberg et al. 2004; Furey et al. 2018), and potentially detrimental to populations (Ydenberg, Tavera, and Lank 2022). However, co-migration can also confer benefits, as interactions may yield useful information, whether inadvertently or intentionally (Danchin et al. 2004; Simons 2004). Research has long demonstrated predator avoidance advantages in flocking birds during daytime (Morse 1970; Pulliam 1973; Lima and Dill 1990). More recently, research found evidence of acoustic communication and interpreted it as a potential benefit during night migration, especially in challenging conditions such as poor visibility or obstacles (Gayk and Mennill 2023; DeSimone et al. 2024; Hüppop and Hilgerloh 2012). In daylight, following conspecifics or heterospecifics en route to breeding grounds may similarly share navigational cues (Couzin et al. 2005). For example, groups of soaring hawks (‘kettles’) may provide social information to multiple species, in this case, about the location of thermal currents that reduce energy expenditure (Leshem and YomTov 1996; Kerlinger and Gauthreaux 1985).

Given migration’s inherent risks (Sillett and Holmes 2002), birds likely leverage available cues — especially from conspecifics — to enhance navigation, speed, and stopover site selection. Yet, the social dimensions of migratory strategies remain underexplored (Schofield et al. 2018; DeSimone et al. 2024). While heterospecific information exchange occurs (Seppänen et al. 2007), it is unclear whether conspecifics provide more useful information than heterospecifics, particularly during active flight. If conspecifics offer distinct advantages, we expect aggregations of the latter within co-migrating flocks. We know conspecific aggregation exists within migrating mixed-species flocks of waterfowl (Eddleman, Patterson, and Knopf 1985; White and James 1978), waders (Erwin 1983; Folmer, Olff, and Piersma 2010) and parrots (Ferdinand et al. 2023). But the presence of conspecific aggregations remains poorly documented in mixed-species flocks of passerines, within or outside migration. Gauthreaux (1972) observed aggregations of conspecifics in 17 species during co-migration in the southeastern United States, but lack of detail prevents a distinction between deliberate and fortuitous clustering. More recently, Gayk et al. (2023) found that temporally adjacent nocturnal migrants in northern Ontario make similar nocturnal flight calls, suggesting conspecific aggregations.

Social interactions between individuals are notoriously difficult to study in the case of co-migrating passerines, because of their tendency to move in loose flocks, mostly at night. However, certain locations along migration routes funnel movements, thus facilitating the observation of possible species aggregations. One such location is the northern coast of the St.Lawrence river estuary, Quebec, Canada (Ibarzabal, Côté, and Drolet 2009). In this location, we tested whether co-migrating passerines formed conspecific aggregations during spring migration events, and whether certain species played a leading role during uninterrupted flights. We looked for conspecific attraction at a fine temporal scale (seconds) and a coarser temporal scale (hours).

## 2 Methods

### 2.1 Study Area

We carried out this study at the dunes of Tadoussac, Québec, Canada (48.157^°^N, 69.665^°^W). This site is on the northern shore of the St. Lawrence river estuary, and is a patchwork of open sandy areas interspersed with deciduous (*Betula sp*., *Populus sp*.) and coniferous (*Pinus sylvestris, Picea glauca*) stands, approximately 50 m above sea level. The dunes of Tadoussac are known as a major hawk migration corridor (Ibarzabal 1999; Brisson-Curadeau, Elliott, and Côté 2020), as well as their spring migration events during which well over 10,000 passerines can sometimes be observed daily, especially near the end of the season (Figure 1). Most of the time, birds fly overhead 10 to 100 m above ground, sometimes they fly over the water, and sometimes birds fly just above ground in a very narrow corridor in the steep slope that borders the estuary. In the spring, flights are generally part of a ‘redetermined’ migration because most birds fly along the coast in a southwest direction, i.e. contrary to their expected direction (Gauthreaux 1978).

**Figure 1.**
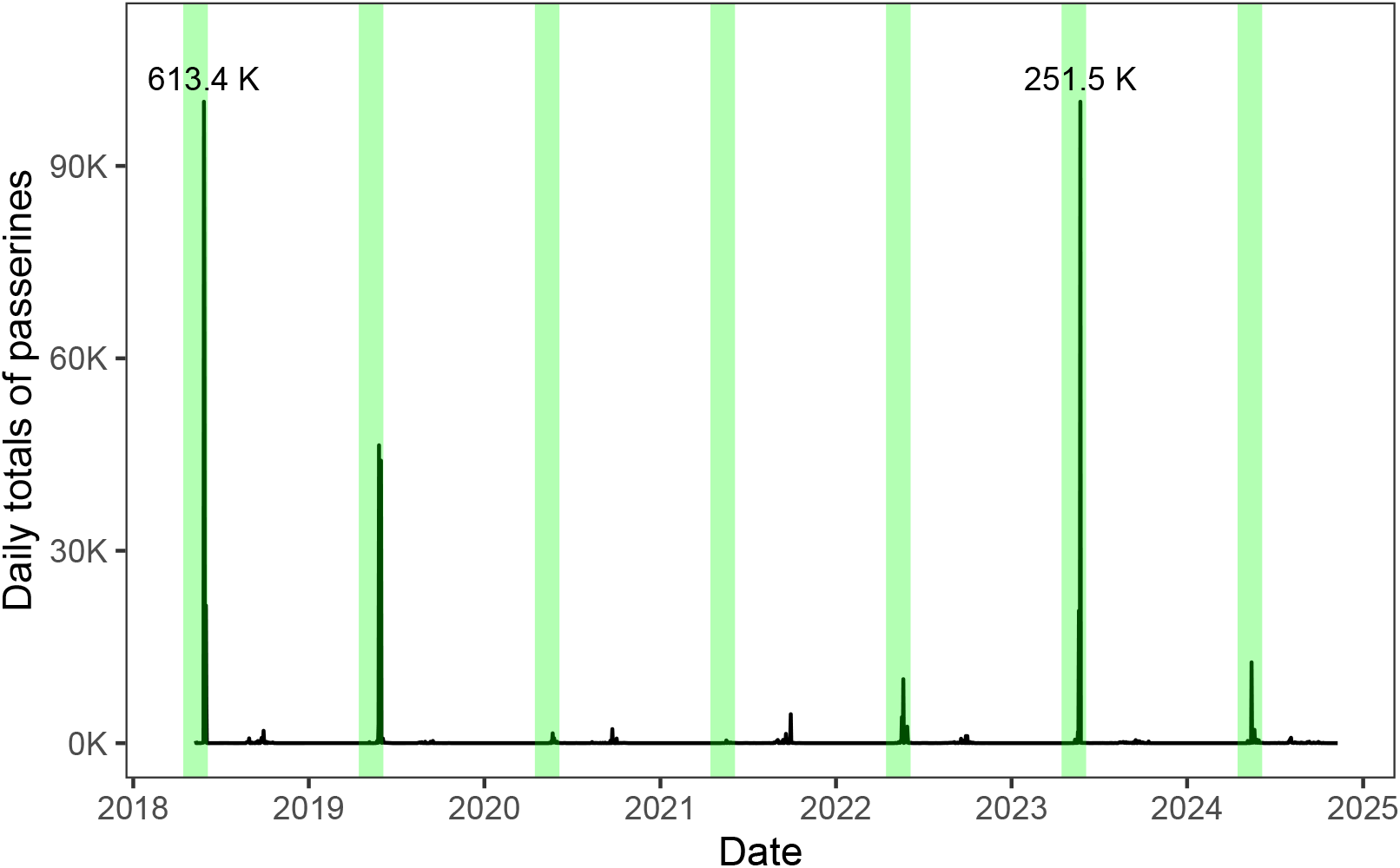
Redetermined migration events at the dunes of Tadoussac, Québec, Canada. Shaded areas are local spring migration seasons for most passerines (15 April - 5 June).

### 2.2 Field methods

We conducted all observations from the top of the slope bordering the estuary. The observation site was situated just above a part of the slope devoid of vegetation, thus facilitating the observation of individual birds, mostly at distances from 10 to 20 m from the observers (Figure 2).

**Figure 2.**
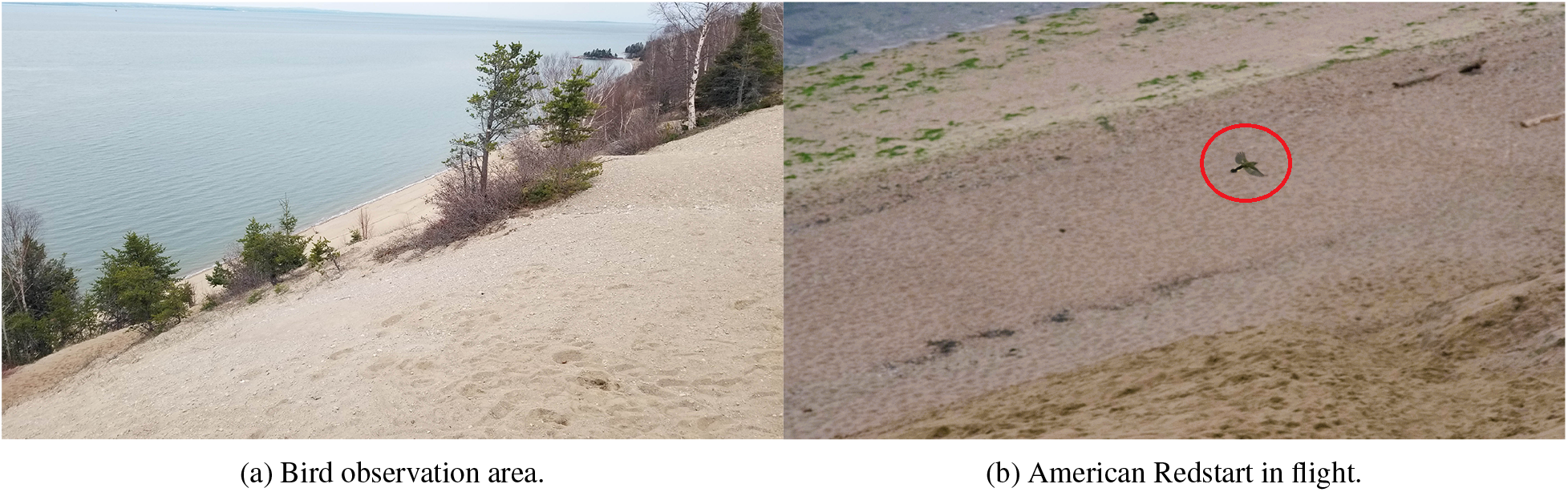
Scenes from the study area.

Redetermined migration events (thereafter, ‘migration events’ or ‘events’) are frequent on the site (Tadoussac Bird Observatory, unpubl. data). Birds during those events often fly ∼ 50 m or higher above ground, in a > 100 m wide corridor. In other cases, they fly hundreds of meters offshore. In the latter cases, most of the individuals are unidentifiable to species. However, when strong winds from north or northwest occur, moving birds tend to fly just above ground in the steep slope facing the estuary, along a ∼ 20 m wide path. This ‘river’ of birds enables the measurement of sequences of individuals identified to species.

We recorded sequences of birds on 12 migration events, each in a separate date, in late May or early June 2019, 2020, 2022, 2024, and 2025. Observations were all performed between 05:00 and 18:05 (EDT) and lasted from 0.8 to 4.6 hours. Each time, two experienced ornithologists recorded sequences of individual birds as they flew from left (NE) to right (SW). One of the ornithologists (coauthor LD) identified each bird, either by naked eye or with binoculars, and named the species or genus out loud. Most species examined here exhibit conspicuous field marks which facilitate migration. Despite this, there was insufficient time in most cases to reliably determine the sex of the bird, and in 12 percent of cases we were unable to identify to species. The second observer (coauthor AD) recorded each bird identification on a smartphone with a CyberTracker (Steventon et al. 2011; Liebenberg et al. 2017) custom-made application showing species acronyms on the screen, requiring just one tap to log species or taxon, with an exact timestamp. This method enabled data collection at rates exceeding 1 bird per second. Despite this, observers were sometimes overwhelmed by large numbers of bird, leading to sequences of unidentified birds.

In addition to the measurement of bird sequences, we performed hourly counts of all birds identifiable to species, either flying low or overhead. We retained only counts from the species observed also during our species sequence sessions (above), but retained counts on all dates, irrespective of migration events.

### 2.3 Statistical analysis

We performed all data preparation and analysis with R/RStudio (R Studio Team 2024; R Core Team 2023). We collapsed all recorded birds into a table with each line representing one individual and its exact passage time. We excluded non-passerine birds, leaving 87783 birds for analysis (Table 1).

**Table 1:** Species and numbers of individuals analysed in redetermined migration events, dunes of Tadoussac, Québec.

| Scientific | Common Name | Code | Number |
| --- | --- | --- | --- |
| *Cardellina canadensis* | Canada Warbler | CAWA | 14 |
| *Cardellina pusilla* | Wilson's Warbler | WIWA | 50 |
| *Catharus fuscescens* | Veery | VEER | 6 |
| *Catharus guttatus* | Hermit Thrush | HETH | 4 |
| *Catharus minimus* | Gray-cheeked Thrush | GCTH | 3 |
| *Catharus ustulatus* | Swainson's Thrush | SWTH | 134 |
| *Empidonax alnorum* | Alder Flycatcher | ALFL | 14 |
| *Empidonax flaviventris* | Yellow-bellied Flycatcher | YBFL | 30 |
| *Empidonax minimus* | Least Flycatcher | LEFL | 37 |
| *Geothlypis philadelphia* | Mourning Warbler | MOWA | 6 |
| *Leiothlypis peregrina* | Tennessee Warbler | TEWA | 482 |
| *Leiothlypis ruficapilla* | Nashville Warbler | NAWA | 29 |
| *Mniotilta varia* | Black-and-white Warbler | BAWW | 13 |
| *Parkesia noveboracensis* | Northern Waterthrush | NOWA | 11 |
| *Seiurus aurocapilla* | Ovenbird | OVEN | 6 |
| *Setophaga americana* | Northern Parula | NOPA | 80 |
| *Setophaga caerulescens* | Black-throated Blue Warbler | BTBW | 42 |
| *Setophaga castanea* | Bay-breasted Warbler | BBWA | 615 |
| *Setophaga coronata* | Yellow-rumped Warbler | YRWA | 3807 |
| *Setophaga fusca* | Blackburnian Warbler | BLBW | 103 |
| *Setophaga magnolia* | Magnolia Warbler | MAWA | 710 |
| *Setophaga palmarum* | Palm Warbler | PAWA | 4 |
| *Setophaga pensylvanica* | Chestnut-sided Warbler | CSWA | 19 |
| *Setophaga ruticilla* | American Redstart | AMRE | 159 |
| *Setophaga striata* | Blackpoll Warbler | BLPW | 44 |
| *Setophaga tigrina* | Cape May Warbler | CMWA | 899 |
| *Setophaga virens* | Black-throated Green Warbler | BTNW | 102 |
| *Vireo olivaceus* | Red-eyed Vireo | REVI | 15 |
| *Vireo philadelphicus* | Philadelphia Vireo | PHVI | 47 |
| *Vireo solitarius* | Blue-headed Vireo | BHVI | 5 |

For each of the migration events, we counted the number of cases where an individual bird was followed by a con-specific. We generated an empirical random distribution of the number of conspecific sequences for each event by reshuffling data 9,999 times. This procedure thus accounts for unequal species counts and missing data. We consid-ered observed counts of conspecific sequences as significantly clustered if their quantile was > 0.995, thus accounting for multiple testing.

To determine if species counts per hour are more clustered than expected under a random (Poisson) distribution, while accounting for varying total counts across hours, we used a Monte Carlo method using Variance-to-Mean Ratios (VMR). We retained all survey dates with at least 5 hourly counts, and one species or mode with at least 10 individuals. For each hour across the days with sufficient data (n = 84), we calculated the observed VMR of counts for each species. We built a null empirical distribution by generating Poisson-distributed counts, accounting for hourly variation of total counts (*λ*) for all species, repeating this 999 times to compute simulated VMRs. We calculated the quantile of the observed VMR in the corresponding null distribution, i.e., the proportion of simulated VMRs that were greater than the observed VMR. We interpreted quantiles > 0.95 as evidence for clustering.

## 3 Results

### 3.1 Sequences of birds

On average, we observed 939 birds per session (range: [113, 3565]). During observation sessions, an average of 6.2 birds/min flowed, but it was highly variable, with long pauses interrupted with intense flows, in some cases exceeding 50 birds/min (Figure 3).

**Figure 3.**
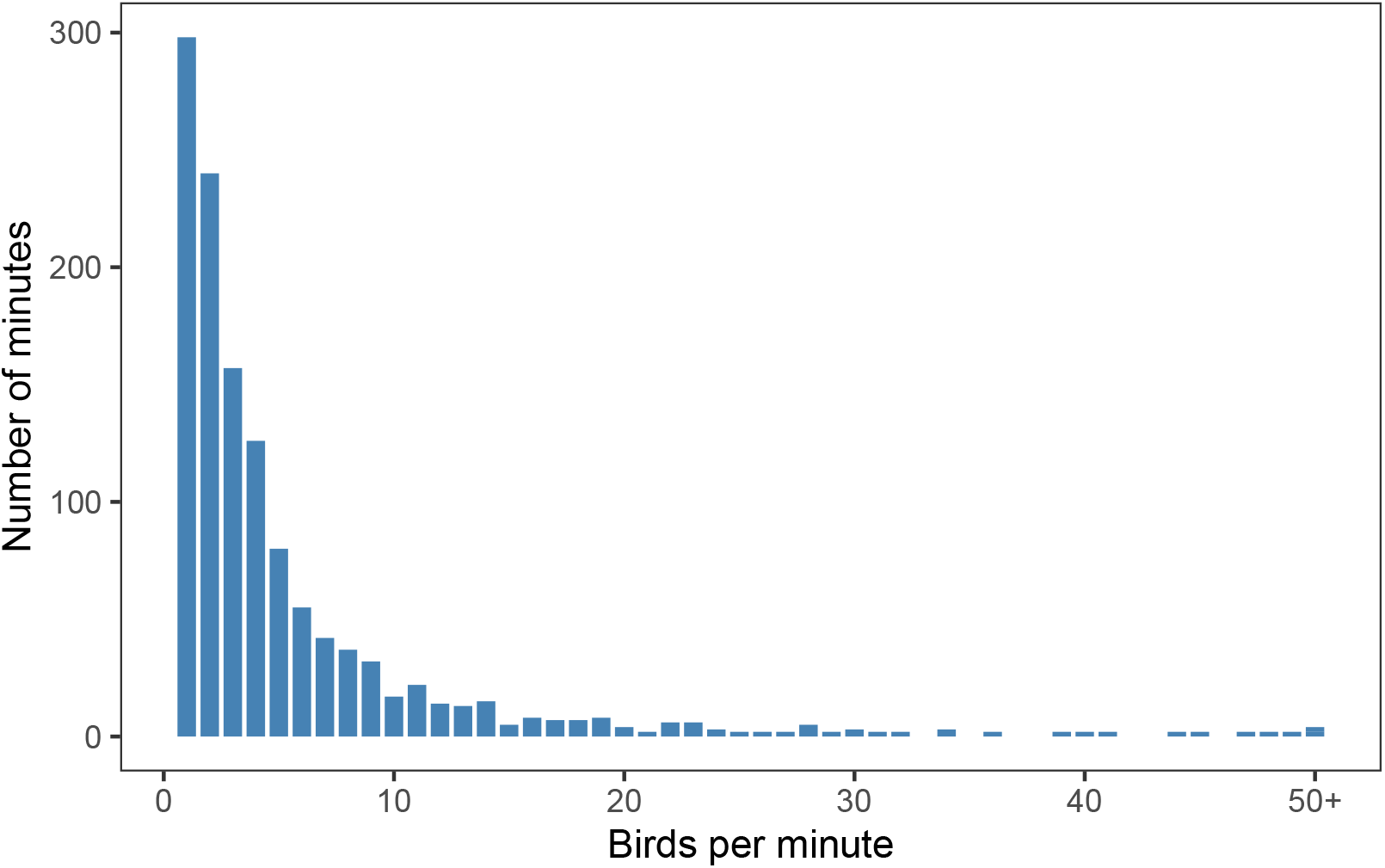
Flow of birds during migration redetermination events, Tadoussac, Québec.

In each migration event, conspecific sequences were significantly more frequent (P < 0.005) than expected if the birds, in the same frequency distribution of species, were mixed independently (Figure 4). Conspecific sequences were often more than 2 birds (max 103 conspecifics in a single run), and the number of 2-bird conspecific sequences (694) was significantly smaller than that expected from a Poisson (random) process (971; *χ*^2^ = 22.9, *P* < 0.001).

**Figure 4.**
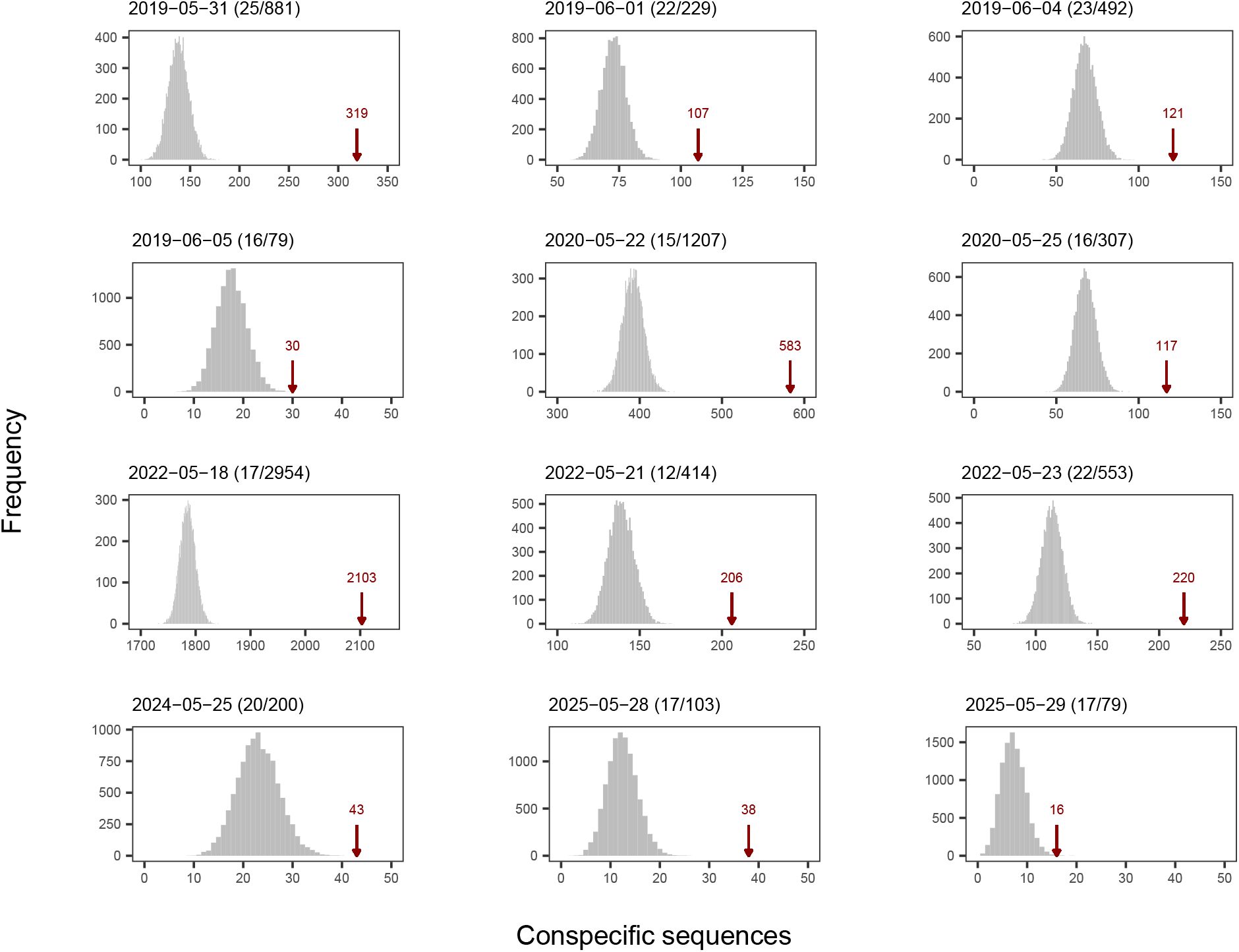
Numbers of conspecific sequences observed vs. expected by chance, on all observation dates. Grey histograms are empirical density functions of conspecific sequences based on the null hypothesis (species independent), and red arrows show actual counts of conspecific sequences. After each date, numbers of species and individuals are shown.

Given the large number of species, interspecific sequences were very variable and most of them occurred a small number of times. However, species sequences that occurred 30 times or more were not asymmetric; that is, there was no evidence that species exerted net leader or follower roles (Table 2).

**Table 2:** Sequences of birds of different species. Only cases with 30+ sequences are shown. with P-values calculated with Fisher’s exact test (two-tailed, null hypothesis of equal frequencies).

| Species 1 | Species 2 | Species 1... |  | P |
| --- | --- | --- | --- | --- |
|  |  | Leading | Following |  |
| American Redstart | Cape May Warbler | 20 | 23 | 0.83 |
| American Redstart | Magnolia Warbler | 30 | 24 | 0.70 |
| American Redstart | Yellow-rumped Warbler | 33 | 24 | 0.45 |
| Bay-breasted Warbler | Cape May Warbler | 68 | 76 | 0.72 |
| Bay-breasted Warbler | Magnolia Warbler | 67 | 61 | 0.80 |
| Bay-breasted Warbler | Tennessee Warbler | 45 | 54 | 0.57 |
| Bay-breasted Warbler | Yellow-rumped Warbler | 165 | 164 | 1.00 |
| Black-throated Blue Warbler | Yellow-rumped Warbler | 19 | 16 | 0.81 |
| Black-throated Green Warbler | Yellow-rumped Warbler | 25 | 18 | 0.52 |
| Blackburnian Warbler | Yellow-rumped Warbler | 27 | 26 | 1.00 |
| Cape May Warbler | Magnolia Warbler | 74 | 70 | 0.91 |
| Cape May Warbler | Tennessee Warbler | 45 | 38 | 0.64 |
| Cape May Warbler | Yellow-rumped Warbler | 278 | 313 | 0.32 |
| Magnolia Warbler | Tennessee Warbler | 32 | 25 | 0.57 |
| Magnolia Warbler | Yellow-rumped Warbler | 176 | 176 | 1.00 |
| Northern Parula | Yellow-rumped Warbler | 20 | 19 | 1.00 |
| Swainson's Thrush | Yellow-rumped Warbler | 26 | 19 | 0.53 |
| Tennessee Warbler | Yellow-rumped Warbler | 116 | 112 | 0.93 |

### 3.2 Hourly clustering

Twenty of the 24 species exhibited significant clustering of hourly counts, at least on some days (Table 3), but with strong day-to-day variation. The tendency for clustering by hours was consistent only for Tennessee Warbler and Magnolia Warbler (Table 3).

**Table 3:** Conspecific clustering by hour, sorted by decreasing order. Days are numbers of dates with at least 10 individuals of the species observed. Clustering is measured as Variance-to-Mean Ratios (VMRs) with their 95 % limits. A VMR of 1.0 would be expected if hourly counts were random (Poisson distributed). Days clustered are numbers of days where a significant clustering occurred. A paired-sampled, one-tailed Wilcoxon test was applied in cases with at least 10 days, to test if species exhibited consistent hourly clustering.

| Species | Days | Variance to Mean Ratio | Days clustered | Wilcoxon P |
| --- | --- | --- | --- | --- |
| Bay-breasted Warbler | 19 | 66.8 [0, 147] | 11 | 0.07 |
| Tennessee Warbler | 41 | 62.2 [0, 135] | 16 | < 0.001 |
| Yellow-rumped Warbler | 75 | 32.1 [12, 52] | 14 | 0.29 |
| Cape May Warbler | 26 | 23.1 [9.6, 37] | 9 | 0.22 |
| Magnolia Warbler | 22 | 19.3 [0, 39] | 9 | 0.03 |
| Blackburnian Warbler | 10 | 19.2 [0, 50] | 5 | 0.22 |
| American Redstart | 8 | 15.8 [0, 37] | 6 |  |
| Northern Parula | 5 | 11.9 [0, 28] | 1 |  |
| Least Flycatcher | 3 | 8.4 [0, 29] | 1 |  |
| Blackpoll Warbler | 3 | 8.2 [0, 33] | 0 |  |
| Black-throated Green Warbler | 11 | 8.2 [1.5, 15] | 3 | 0.21 |
| Swainson's Thrush | 9 | 7.9 [4.4, 11] | 6 |  |
| Wilson's Warbler | 4 | 7.8 [0, 19] | 2 |  |
| Red-eyed Vireo | 1 | 6.7 | 1 |  |
| Yellow-bellied Flycatcher | 1 | 6.2 | 1 |  |
| Philadelphia Vireo | 3 | 5.7 [0, 14] | 2 |  |
| Nashville Warbler | 8 | 4.8 [1, 9] | 2 |  |
| Chestnut-sided Warbler | 4 | 4.6 [0, 9] | 2 |  |
| Black-throated Blue Warbler | 5 | 4.5 [0, 10] | 1 |  |
| Orange-crowned Warbler | 1 | 4.3 | 1 |  |
| Mourning Warbler | 1 | 3.4 | 1 |  |
| Northern Waterthrush | 2 | 2.5 [0, 14] | 0 |  |
| Black-and-white Warbler | 2 | 2.5 [1.4, 4] | 0 |  |
| Canada Warbler | 3 | 2.4 [1.2, 4] | 0 |  |

## 4 Discussion

Given that most migratory songbirds exhibit species-specific peak migration dates, conspecific clustering at coarse temporal scales (e.g., weeks) is expected. But to our knowledge, this is the first detailed evidence of conspecific attraction at finer temporal scales, hours and seconds, during migration. On given days, overall migration intensity usually varied markedly among hours, but we found that single-species numbers often varied even more, providing evidence for conspecific attraction at the temporal resolution of hours. At a finer, behavioral, resolution of seconds, conspecific attraction was evident, and occurred on each opportunity we had to observed precise sequences of birds actively flying.

Most of the literature on conspecific attraction has focused on the breeding season, following the pioneering study by Stamps (1988). While poorly documented, conspecific attraction during migration has often been inferred from casual or indirect observations. Gauthreaux speculated that conspecifics possibly gather in flocks at dawn before completion of their solo flight across the Gulf of Mexico (Gauthreaux 1972). Balcomb also believed that most passerine birds flew singly at night (Balcomb 1977). An early study in Louisiana suggested the occurrence of conspecific attraction during landfall at dawn (Gauthreaux 1972). Hamilton (1962) and Ball (1952) also came to the conclusion of conspecific attraction triggered or maintained by calls. A more recent study (Gayk and Mennill 2023) documented spatial aggregation of similar nocturnal flight calls in Ontario, again suggesting that passerines of the same species flock together during nocturnal migration.

As for the hourly clustering, its sporadic occurrence in most species could reflect opportunistic responses to temporary cues like food, weather, or predation risk. The consistent clustering in Tennessee and Magnolia and possibly, Bay-breasted Warblers might indicate stronger social drivers.

Our evidence for conspecific attraction at the scales of seconds and hours is novel, and before before attempting to place it in a conceptual frame, replication will be needed. For now, we can only speculate on the mechanisms leading to those patterns. One plausible explanation would be the facilitation of information sharing among individuals with similar ecological needs (Couzin et al. 2005; Leshem and YomTov 1996; Kerlinger and Gauthreaux 1985). For instance, collective decision-making in conspecific flocks could enhance the detection of favorable winds or food patches, as suggested by studies on collective navigation in migratory birds (Simons 2004). In contrast, heterospecific flocks might introduce conflicting cues due to differing migratory strategies or destinations, reducing the reliability of group-level decisions.

From an evolutionary perspective, conspecific attraction during migration may also reflect selection pressures related to territory acquisition and more generally, breeding preparation. Migratory songbirds, particularly males, often arrive at breeding grounds earlier than females - a phenomenon known as protandry (Morbey and Ydenberg 2001) — and establishing conspecific networks en route could facilitate territory defense or mate attraction upon arrival. Flying with conspecifics might allow individuals to assess competitors or form alliances.

The preference for conspecific over heterospecific associations contrasts with some evidence of mixed-species flocks during migration, particularly at stopover sites (Goodale et al. 2010). These heterospecific groupings are often attributed to predator avoidance or resource competition, benefits that may be more pronounced during stationary phases than active flight. Our findings, focused on in-flight behavior, suggest that the costs of coordinating with heterospecifics—such as mismatched flight speeds or navigational goals—may outweigh these advantages during transit.

Several limitations in our study should be addressed in future research. First, we were unable to determine the sex of most birds, including those with sexually dimorphic plumages thus precluding inference about birds flying in pairs, or comparisons of conspecific attraction within sex. Second, we can only speculate whether conspecific attraction was driven by navigation, communication, or social mechanisms. Experiments manipulating flock composition or simulating conspecific cues could clarify these drivers.

We were unable to follow individual birds for more than a few seconds, thus we cannot determine how ephemeral are species aggregations. It is possible that some individuals fly with their mate, which could explain the tendency of conspecifics to be together. However, two observations point against this explanation. First, there were not more sequences of two conspecifics than expected by chance - in fact there were significantly fewer. Furthermore, as noted earlier, males usually migrate sooner in the season than females in many passerine species (Morbey and Ydenberg 2001).

In conclusion, our finding that songbirds preferentially fly with conspecifics during migration underscores the importance of species-specific sociality in shaping migratory behavior. This conspecific attraction likely enhances navigational efficiency, reinforces communication, and prepares individuals for breeding, offering a selective advantage in a challenging life-history stage. These insights contribute to our understanding of social dynamics in migratory birds and open avenues for exploring how such behaviors influence population connectivity and resilience. As migratory songbirds face increasing threats from habitat loss and climate change, unraveling these social strategies may also inform conservation efforts, ensuring that critical group behaviors are preserved across their annual cycle.

## 5 Acknowledgments

We thank birders at the Observatoire d’oiseaux de Tadoussac, who not only provided help by finding birds, but also entertainment from lively discussions in those frequent moments when no birds are to be found. Thanks especially to Alexandre Terrigeol, Pascal Côté, and Jean-François Therrien, for comments on earlier versions of this manuscript. Funding for this study was provided by Université Laval department funds, and Explos-Nature.

